# Absolute and Relative Declines in National Institutes of Health (NIH) Funded Basic Science Publications

**DOI:** 10.1101/2025.07.15.664939

**Authors:** Michael S Lauer

## Abstract

An analysis of over 2.2 million NIH-funded research papers published between 1990 and 2024 reveals that there have been marked declines in the relative and absolute numbers of basic-science oriented publications. In 1990, nearly 60% of NIH-funded publications reported on basic science, with that proportion declining to 24% in 2024. The absolute number of basic science publications increased after the NIH budget doubling of 1998-2023, but at a slower rate than for human-focused publications. After the 2011-2013 budget shocks, the absolute number of basic science publications declined. These data support concerns about the long-term trajectory of NIH’s support for basic science.

## Introduction

When he retired in 2025, former Director Francis Collins stated that that the National Institutes of Health (NIH) is “rightfully called the crown jewel of the federal government for decades.”(1) NIH-funded science led to the development of nearly all new drugs approved by the Food and Drug Administration from 2010-2099 and to over 100 Nobel prizes.(2) Most of the science that has led to new drugs and Nobel prizes has been basic, or fundamental, science, which the US government defines as “research directed toward increasing knowledge a fuller knowledgeor understanding of the subject under study, rather than any practical application of that knowledge.”(3)

Thought leaders inside (4, 5) and outside (6) NIH have expressed concern that the agency has excessively de-emphasized basic science in its quest to enhance translation and fund clinical research. In 2020, Dr. Jay Battacharya, now NIH Director, described decreasing agency support for “edge science,” which most often is basic science.(7) NIH conducted its own textual analysis of over 190,000 parent awards and linked subprojects funded between 2009 and 2022 and reported a marked decrease in support for basic science, with decreasing number of projects and amount of funds.(8) Expressing concern, Senator Bill Cassidy (R-LA) wrote, “Waning federal focus on basic research could lead to a decline in treatments and cures eventually developed through private funding. Unlike certain areas of clinical research, the private sector would not be equipped to fill gaps in support for basic research.”(9)

Aside from analyzing grants, another approach to assessing secular portfolio trends leverages the National Library of Medicine’s Medical Subject Heading (MeSH) taxonomy of publications. Weber described a “triangle of biomedicine” that relies on MeSH terms for human, animal, and molecular/cellular science to classify different types of science.(10) The NIH Office of Portfolio Analysis (OPA) adapted and slightly modified Weber’s triangle to create a publicly available tool by which publications can be classified as 1) human-focused, meaning that there are human MeSH terms but no animal or molecular/cellular terms, 2) fundamental, meaning that there are animal and/or molecular/cellular terms but no human terms, or 3) mixed.(11) Here I describe an analysis of NIH-funded papers published between 1990 and 2024. I find that there have been marked relative and absolute declines in NIH-supported basic science publications which are coincident with exogenous budget shocks and certain policy changes.

## Results

I found 2,214,497 NIH-funded research publications, of which 823,455 (37%) were fundamental and 760,264 (34%) were human focused. Among the papers with human as well animal and/or molecular/cellular MeSH terms, 427,380 (19%) had more molecular/cellular terms (i.e., mixed mostly fundamental) and 203,398 (9%) had more human terms (i.e., mixed mostly human).

In the 1990s, most NIH publications reported fundamental science or mixed mostly fundamental science (Figure 1A). After the NIH budget doubling (1998-2003) the number of NIH publications increased, but to a greater degree among human-focused publications, which exceeded all other types by 2012. After the passage of the Budget Control Act of 2011 and imposition of budget sequestration in 2013, the number of fundamental and mixed mostly fundamental papers declined and the number of human-focused papers plateaued. With the COVID-19 pandemic in 2020, the number of human-focused and mixed mostly human publications increased, while the number of fundamental publications plateaued.

**Figure 1.**
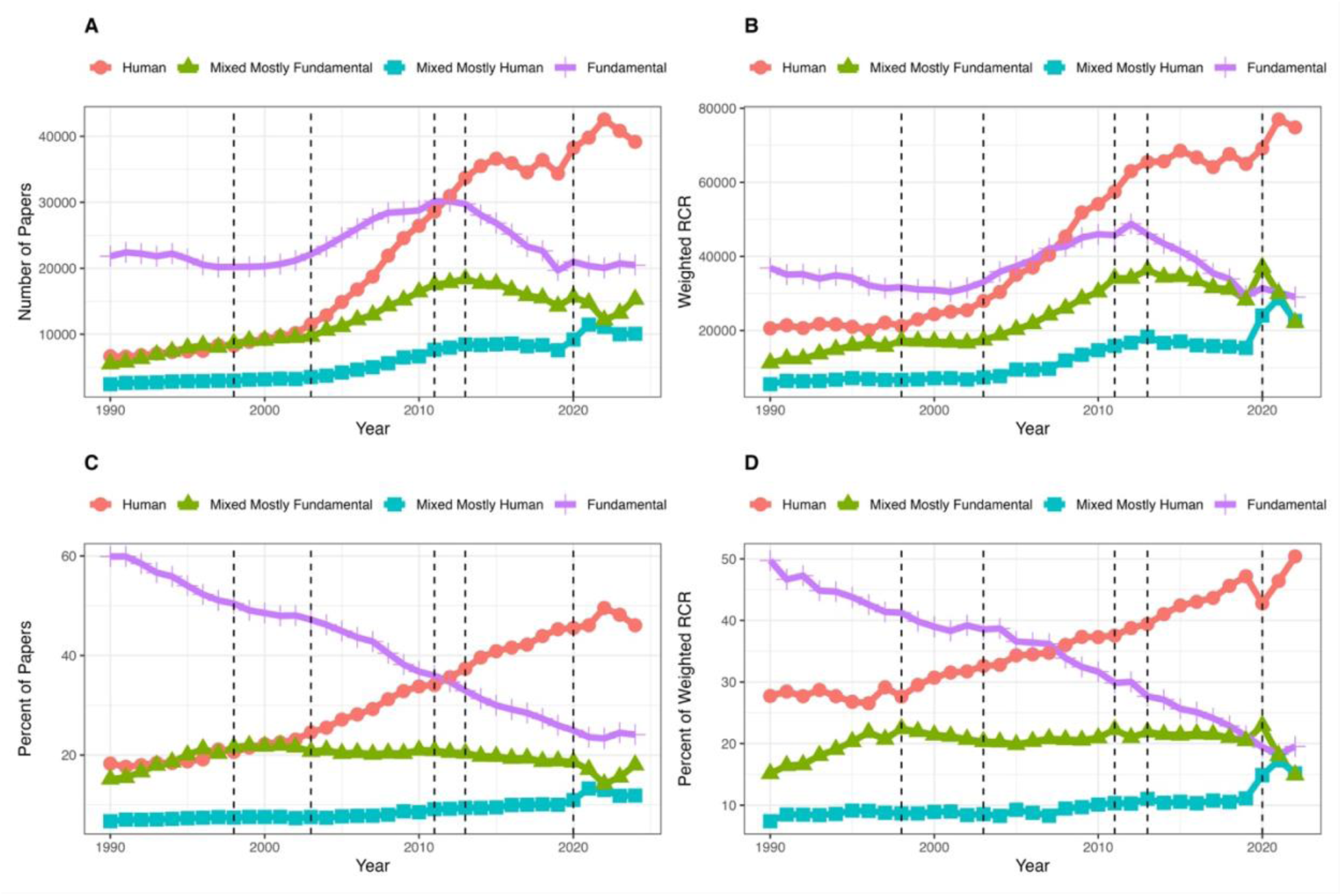
Secular trends in the types of science reported in NIH-funded publications. Based on National Library of Medicine Medical Subject Heading (MeSH) terms, publications are classified as human-focused, fundamental, mixed mostly fundamental, and mixed mostly human. Vertical lines refer to the NIH budget doubling from 1998 to 2003, the 2011 Budget Control Act which led to budget sequestration in 2013, and the COVID-19 pandemic in 2020. Panel A shows absolute numbers of papers. Panel B shows numbers of papers weighted for each paper’s relative citation ratio (RCR). Panel C shows the proportion of papers in each category. Panel D shows the proportion of papers in each category after weighting each paper for its relative citation ratio.

Not all publications have equal influence, so I repeated the analysis weighting each publication by its relative citation ratio (RCR), a bibliometric measure developed by the NIH OPA which normalizes citation numbers for field and year of publication.(12) An RCR of one implies that a paper received the median number of citations for NIH-funded papers in the same field and published in the same year. If a publication garners no citations its RCR-weighted value is zero, while if it garners 100 times more citations than the NIH field-specific median its RCR-weighted value is 100. I plotted RCR-weighted values for different types of papers and found similar patterns (Figure 1B), though there was a marked peak for mixed mostly human papers concurrent with the COVID-19 pandemic.

The relative decline in fundamental science publications dates back to at least 1990, when fundamental and mixed mostly fundamental science accounted for 75% of publications and human-focused science accounted for only 18% (Figure 1C). In 2024, human-focused and mixed mostly human science accounted for 58% of publications, while fundamental and mixed mostly fundamental science accounted for only 42%. The long-term patterns were similar after I weighted each paper for its relative citation ratio (Figure 1D).

These trends may be reflective of more general trends, trends that transcend the NIH ecosystem. Therefore I obtained PubMed identification numbers for all non-NIH funded research papers published between 1990 and 2024 and used the OPA’s tool to classify a 2% random sample for each year according to type of science. I used the same classification scheme as I did for NIH funded research: human-focused, fundamental, mixed mostly fundamental, and mixed mostly human. Secular trends for non-NIH papers were distinctly different (Figure 2). During all years, most papers were human-focused, and the relative proportions remained fairly constant.

**Figure 2.**
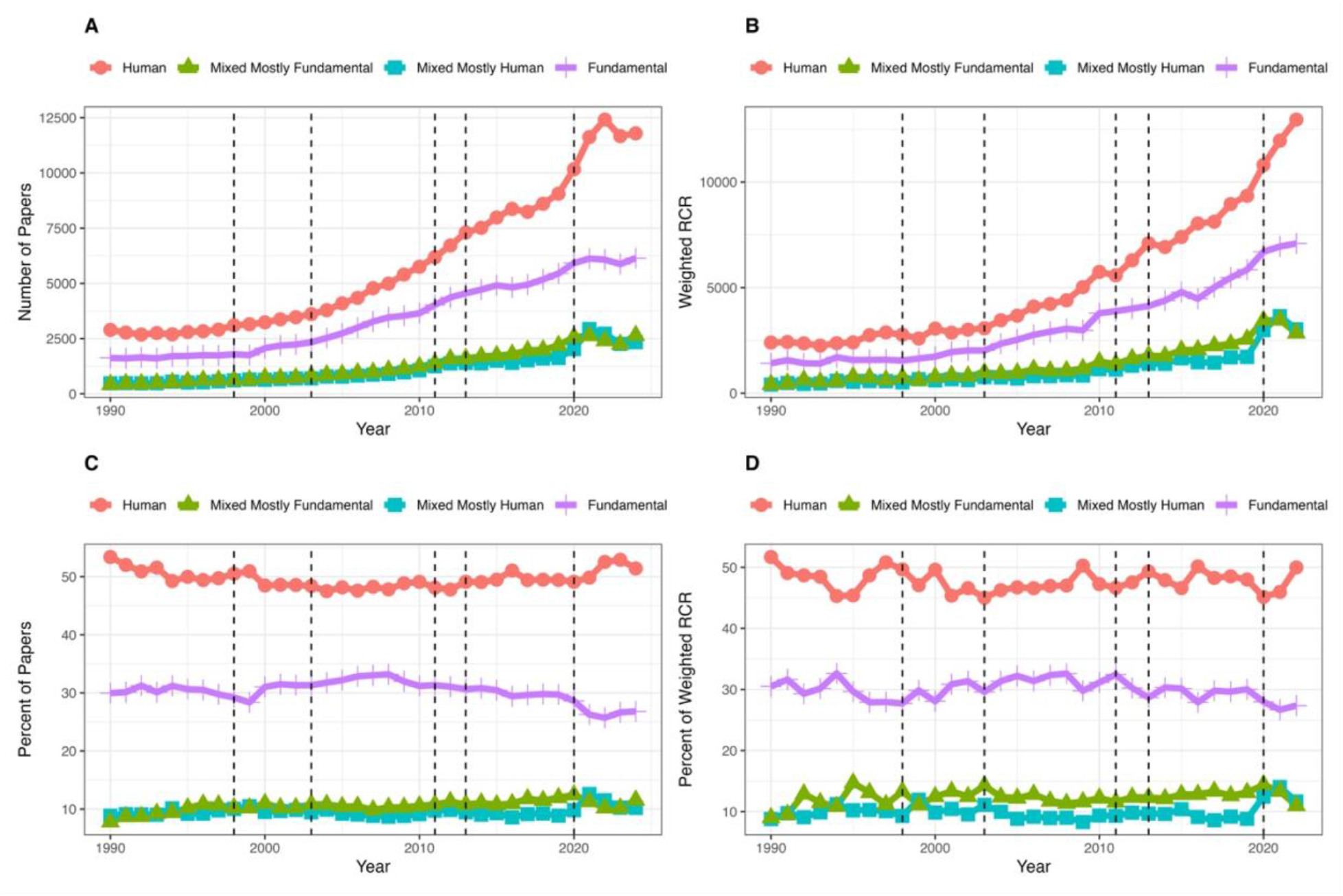
Similar data as shown in Figure 1 except for non-NIH-funded publications. Data are based on a 2% random sample of all PubMed research papers. Panels are analogous to those shown in Figure 1.

The different trajectories for NIH-funded and non-NIH funded papers are clear when shown in Weber’s triangle of biomedicine (Figure 3). In the 1990s, NIH-funded papers remained stable, moving slightly away from animal and toward molecular/cellular science. In the 2000s and 2010s there were marked shifts toward human-focused science, away from fundamental science.

**Figure 3.**
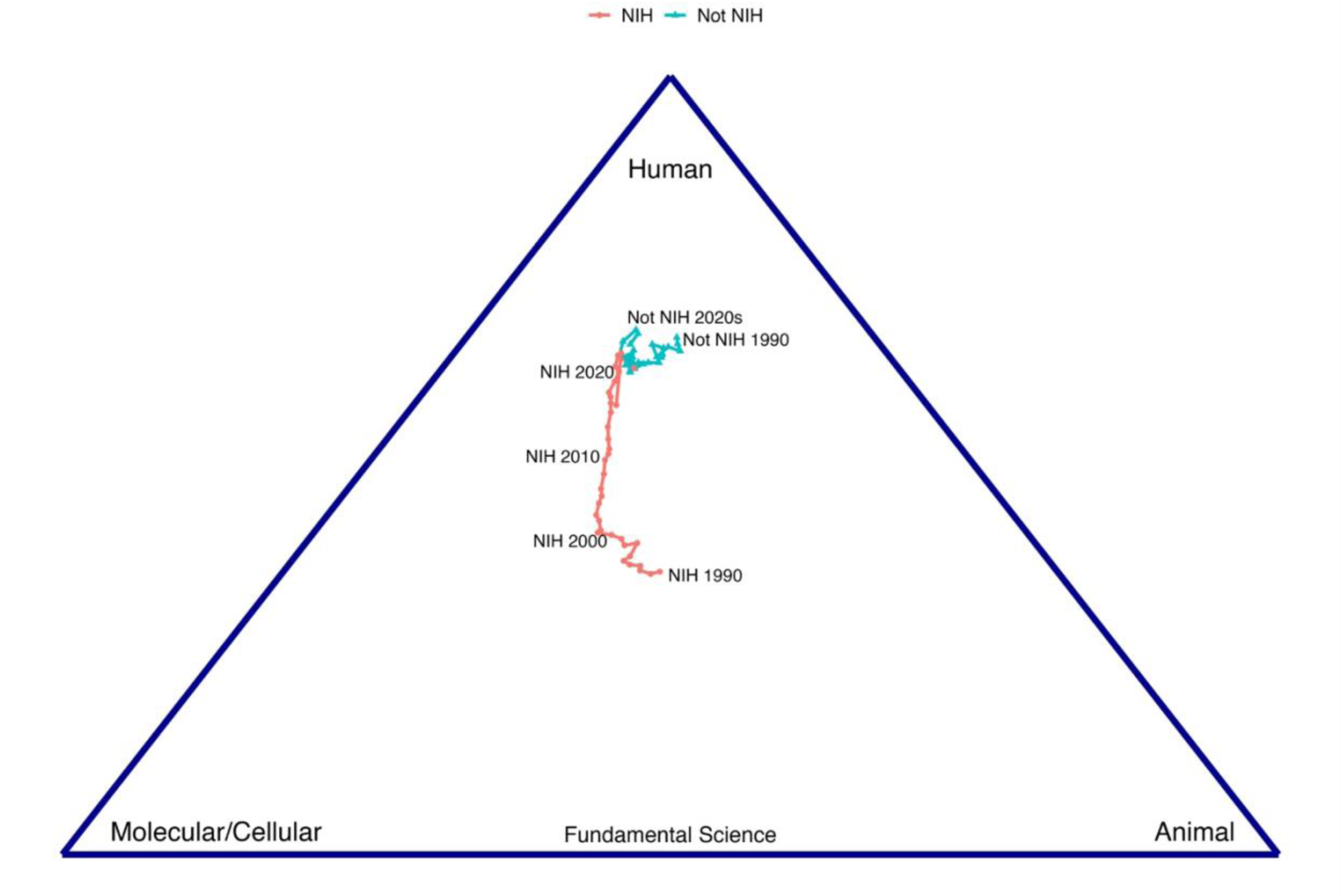
Secular trends in the NIH-funded and non-NIH funded research papers according to Weber’s triangle of biomedicine (references 10 and 11). Each point represents the average coordinates for papers published each year proceeding from 1990 to 2024.

## Discussion

These data support concerns about the long-term trajectory of NIH’s support for basic science.(4–6) In the 1990s, NIH-supported scientists largely published on fundamental science. Marked changes occurred with the NIH budget doubling of 1998-2003, the establishment of the NIH Roadmap for Medical Research in the early to mid 2000s, the establishment and growth of the Clinical Translational Science Award (CTSA) program from 2006-2012, and the budget shocks following the passage of the 2011 Budget Control Act. There were changes with the COVID-19 pandemic, but whether these changes will affect long-term trends is unclear. NIH now appears to be facing major cuts in budget support, which may have further implications for the agency’s ability to support basic science.(13)

## Material and Methods

I obtained PubMed ID numbers for NIH-supported publications from NIH ExPORTER linkage files (14) and for non-NIH supported publications from PubMed through their EDirect system.(15) I used the NIH OPA’s iCite tool (11) to classify papers as fundamental, human-focused, or mixed and to obtain each paper’s RCR.

## Appendix Additional information on methods and data

### Data sources

I obtained NIH publication data from annual files posted on NIH’s ExPORTER site (https://reporter.nih.gov/exporter/publications). I identified unique PubMed ID numbers (PMIDs) and fed them in aliquots of at most 50,000 into the NIH Office of Portfolio Analysis’s (OPA) iCite tool (https://icite.od.nih.gov/analysis). The resulting output contains values for “human,” “animal,” and “molecular/cellular,” with each referring to a class of MeSH terms and each having a value of 0 to 1 depending on what proportion of all MeSH terms fit into that category. Thus, if a publication has 9 human MeSH terms and 1 molecular/cellular term, the values for human, animal, and molecular/cellular would be 0.9, 0, and 0.1. If a publication has no human, animal, or molecular/cellular MeSH terms, each has a value of 0; this occurs with < 5% of NIH-funded publications. Otherwise, the values must add up to one.

The iCite tool also returns each paper’s relative citation ratio (RCR) and a determination as to whether the paper reports on research findings, as opposed to a review or commentary.

I limited all analyses to papers that had at least one human, animal, or molecular/cellular MeSH term and that the iCite tool classified as a research article.

I obtained PMIDs for non-NIH research papers by working within a Unix environment and interrogating PubMed’s EDirect tool (https://hpc.nih.gov/apps/edirect.html). For each year my command line was:

esearch -db pubmed -query ‘YEAR[PDAT] NOT Review[pt] NOT NIH[gr] NOT Book[pt] NOT Letter[pt] NOT Editorial[pt] NOT News[pt] NOT Comment[pt]’ | efetch -format uid > all_pmids_YEAR.txt

I used the Hadley Wickham’s R tidyverse package’s (https://www.tidyverse.org/) slice_sample command to pull out a 2% random sample for each year (i.e., 2% of 1990 PMIDs, 2% of 1991 PMIDs, through to 2% of 2024 PMIDs). I ran each of these samples through iCite in the same way that I did for the NIH-funded PMIDs, and again limited all analyses to papers that had at least one human, animal, or molecular/cellular MeSH term and which the iCite tool classified as a research article.

### Classification

I illustrate in the table below how I classified each paper; my approach follows B.

I. Hutchins, M. T. Davis, R. A. Meseroll, G. M. Santangelo, Predicting translational progress in biomedical research. *PLoS Biol* **17**, e3000416 (2019).

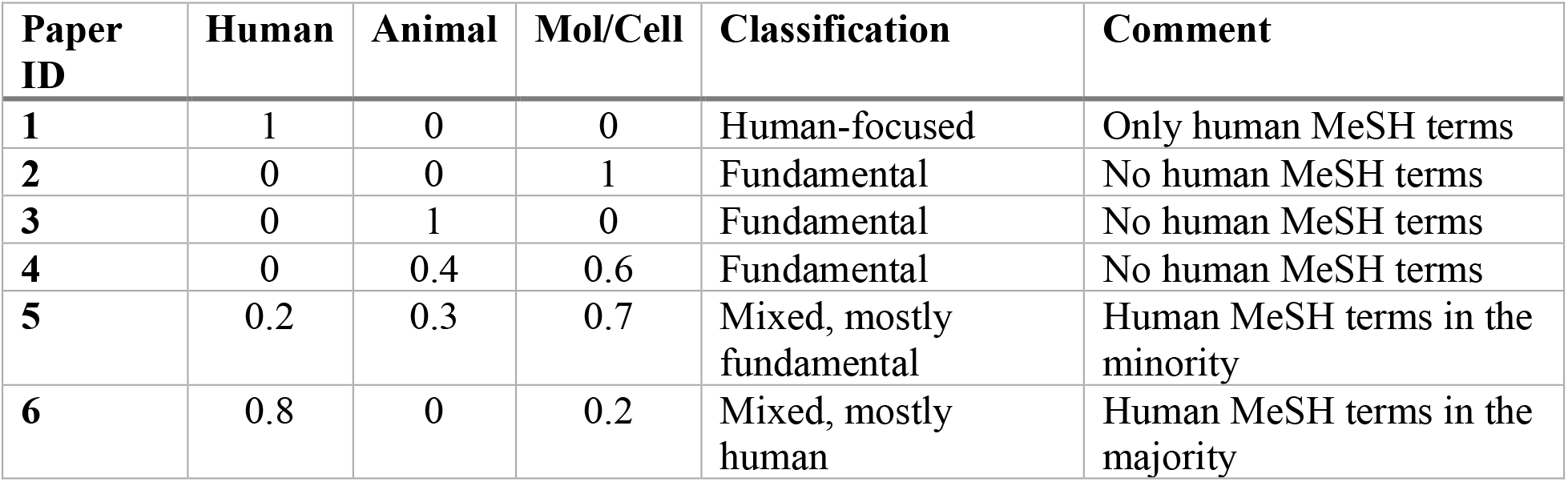

### Plots

I made all figures with Wickham’s ggplot2 package (https://ggplot2.tidyverse.org/).

### Sharing

I will make all data frames and code available when the paper is accepted for publication.

## References

1. Dr. Francis Collins retires from NIH, urging support for crucial research and embattled workers. PBS News (2025). Available at: https://www.pbs.org/newshour/politics/dr-francis-collins-retires-from-nih-urging-support-for-crucial-research-and-embattled-workers [Accessed 10 July 2025].

2. Long a ‘Crown Jewel’ of Government, N.I.H. Is Now a Target - The New York Times. Available at: https://www.nytimes.com/2024/12/01/health/nih-trump-kennedy-bhattacharya.html [Accessed 10 July 2025].

3. Definitions of Research and Development: An Annotated Compilation of Official Sources | NSF - National Science Foundation. Available at: https://ncses.nsf.gov/pubs/ncses22209#iii-federal-and-state-government-r-d_b-federal-acquisitions-regulations_definition [Accessed 10 July 2025].

4. J. Kaiser, Neurological Institute Finds Worrisome Drop in Basic Research. Science (1979) (2014). 10.1126/article.23319.

5. J. R. Lorsch, L. A. Tabak, M. M. Bertagnolli, Applied research won’t flourish withoutbasic science. Elife 13 (2024).

6. M. Rosbash, A Threat to Medical Innovation. Science (1979) 333, 136–136 (2011).

7. M. Packalen, J. Bhattacharya, NIH funding and the pursuit of edge science. Proceedings of the National Academy of Sciences 117, 12011–12016 (2020).

8. M. Lauer, Trends in NIH-Supported Basic, Translational, and Clinical Research: FYs 2009-2022 – NIH Extramural Nexus. Available at: https://web.archive.org/web/20241031125926/ https://nexus.od.nih.gov/all/2023/10/31/trends-in-nih-supported-basic-translational-and-clinical-research-fys-2009-2022/ [Accessed 10 July 2025].

9. NIH in the 21st Century: Ensuring Transparency and American Biomedical Leadership.

10. G. M. Weber, Identifying translational science within the triangle of biomedicine. J Transl Med 11, 126 (2013).

11. B. I. Hutchins, M. T. Davis, R. A. Meseroll, G. M. Santangelo, Predicting translational progress in biomedical research. PLoS Biol 17, e3000416 (2019).

12. B. I. Hutchins, X. Yuan, J. M. Anderson, G. M. Santangelo, Relative Citation Ratio (RCR): A New Metric That Uses Citation Rates to Measure Influence at the Article Level. PLoS Biol 14, e1002541 (2016).

13. Trump’s proposed budget details drastic cuts to biomedical research and global health. Science (1979) (2025). 10.1126/SCIENCE.ZE5W9SI.

14. NIH ExPORTER. Available at: https://reporter.nih.gov/exporter/publications [Accessed 10 July 2025].

15. Entrez Direct E-utilites. Available at: https://hpc.nih.gov/apps/edirect.html [Accessed 10 July 2025].

